# Bacterial secreted products selectively inhibit non-symbiotic fungi in bees

**DOI:** 10.64898/2026.09.16.752052

**Authors:** Lilian Caesar, Amadeus Wagner, Gabriela Toninato de Paula, Monica T. Pupo, Irene Newton

## Abstract

Microbial interactions play an important role in shaping microbiome assembly, such as by limiting invasion by harmful organisms that can directly affect the host or disrupt microbiome-associated benefits. Such interactions have been observed across systems, including in the microbiomes of key pollinators such as stingless bees. In *Scaptotrigona depilis*, bacteria associated with the larval diet inhibit potentially pathogenic filamentous fungi while allowing beneficial yeast symbionts to persist. The mechanisms underlying these effects, however, remain unclear. Here, we combined conditioned media (cell-free supernatant) assays with genomic and metabolomic analyses to investigate whether bacterial secreted products mediate these effects in the bee microbiome. Our results show that bacterial secreted products, particularly from prevalent bacterial taxa such as *Apilactobacillus kunkeei*, strongly affect fungal growth. Filamentous fungi, including the pathogen *Aspergillus*, were consistently inhibited, partly through substrate acidification driven by organic acids, but also through additional acidity-independent factors. In contrast, yeast responses were more variable: a non-symbiotic *Zygosaccharomyces* was inhibited by bacterial metabolites under near-neutral pH, whereas the symbiotic *Zygosaccharomyces* required for larval development was maintained or promoted under acid-conditioned media. Genomic analyses revealed limited canonical antifungal biosynthetic clusters in the most prevalent bacteria in the larval diet, while metabolomics identified extracellular peptide-like compounds across strains, suggesting a role for non-canonical secreted products. Together, these results show that bacterial secreted products play a key role in selectively shaping fungal communities in the stingless bee larval diet, providing first hints on a mechanistic basis for how microbial interactions structure this ecosystem.

**Importance:** Microbes associated with hosts or other systems have evolved strategies to persist in their environments, including inhibiting invading microorganisms that could harm their hosts or disrupt the microbial community. In systems where multiple microbial partners coexist, however, these interactions must be selective. The stingless bee *Scaptotrigona depilis*, for example, hosts bacteria and fungi in its larval diet, where bacteria inhibit fungi outside the core community while allowing required yeast symbionts to persist. Here, we show that this selective fungal inhibition can be mediated by substances released by bacteria, without requiring direct cell-to-cell competition. Bacteria modify the extracellular environment and inhibit potentially harmful fungi, including *Aspergillus*, through acidification and additional secreted products, while allowing the beneficial yeast to persist. These findings provide a mechanistic basis for selective bacterial-fungal interactions and point to bacterial secreted products as a potential source of yet-uncharacterized molecules with applications in microbial control.

## Introduction

Microbiomes play important roles in host health not only through direct interactions with the host, such as nutrient provisioning and immune modulation, but also indirectly by shaping microbial community structure (1). Interactions among microbes contribute to determine which taxa persist, and this is essential for maintaining community function. In many systems, microbiomes or specific members of these communities limit invasion by harmful strains, thereby protecting both the resident community and its associated benefits (2–4). The mechanisms underlying these interactions vary across systems and resolving them is essential to linking microbiome function to host outcomes, as well as informing applications to health, conservation, and bioactive compound discovery.

Eusocial bees represent a particularly relevant system to investigate such mechanisms. These key pollinators harbor stable microbiomes that contribute to their health (1), yet they are continuously exposed to environmental microbes that threaten their homeostasis. During foraging, worker bees interacting with plants and pollinators can acquire diverse microorganisms, which they subsequently bring back to the colony, introducing potentially competing or pathogenic taxa (5, 6). Because bees live in densely populated societies with frequent social contact, this exposure increases the risk of disease transmission even more (7, 8). As a result, microbial contamination in any part of the colony—from adult individuals to food stores and developing larvae and their diet—can have significant consequences for colony stability and survival.

Fungi represent an important class of microbial threats in bee colonies, particularly affecting developing larvae and contributing to colony losses (9). In honey bees, well-characterized fungal pathogens include *Ascosphaera apis*, the causative agent of chalkbrood, and *Aspergillus* spp., responsible for stonebrood. As in other host-associated systems, the bee microbiome has been shown to play a key role in limiting such infections. For example, *Apilactobacillus kunkeei*, commonly associated with bee brood, and *Bifidobacterium* spp., prevalent in the adult gut, can inhibit *A. apis* growth *in vitro* (10, 11). Similarly, the larva-associated bacterium *Bombella apis* inhibits *Aspergillus* sp. growth and sporulation both *in vivo* and *in vitro* (2). In stingless bees, the only other group of highly eusocial bees forming large perennial colonies (12), some studies have also reported antifungal activity by associated bacteria. For instance, *Levilactobacillus*, *Acetobacter*, *Lactiplantibacillus*, and *Pantoea* isolated from stingless bee workers can inhibit *A. apis* and *Aspergillus flavus* (13).

Bacterial inhibition of fungi can be mediated by multiple mechanisms, including nutrient depletion, spatial competition, and the production of extracellular inhibitory compounds (14). Among these, secreted compounds are frequently implicated, as suggested for bee-associated bacteria that inhibit the growth of *A. apis* and *Aspergillus* (2, 10, 11, 13). Conditioned media (cell-free supernatant) assays have shown that bacterial secreted products alone can be sufficient to inhibit fungal growth, highlighting the potential role of compounds such as organic acids, peptides, and specialized metabolites, often encoded by biosynthetic gene clusters (BGCs). However, only one study has followed up on these observations to identify candidate compounds or underlying mechanisms. In *Bombella apis*, genome annotation identified biosynthetic gene clusters, including a type I polyketide synthase (T1PKS), which has been proposed as a candidate contributor to its antifungal activity (2), given the known association of T1PKS-derived compounds with antifungal functions (15). In other cases, mechanisms or secreted products have not been investigated, and for several taxa there is limited information on overall ecological relevance within the studied bee colonies, such as the prevalence or distribution of these microbes within the colony (13).

By combining microbiome characterization of a stingless bee colony with *in vitro* competition assays between bacterial and fungal isolates, we previously identified a novel case of bacterial symbionts exhibiting protective antifungal activity (16). In *Scaptotrigona depilis*, the larval diet hosts both transient, potentially pathogenic fungi and a core yeast community (16), including *Zygosaccharomyces*, which is required for larval development (17), making it a particularly informative system to study bacterial-fungal interactions. We found that *Apilactobacillus*, a representative strain of abundant taxa in the larval diet belonging to the *Apilactobacillus-Acetilactobacillus-Nicoliella* clade, strongly inhibits potentially pathogenic fungi while promoting the symbiotic yeast (16). Other isolates from the same hive site, including *Weissella* and *Bombella*, also exhibited antifungal activity, although these effects were weaker or less selective. However, these assays relied on direct competition and did not identify the potential mechanisms underlying these interactions.

Here, we tested whether the effects of bacteria on fungal growth previously observed for members of the *Scaptotrigona depilis* microbiome are mediated by bacterial secreted products. Using representative bacterial isolates (*Apilactobacillus*, *Weissella*, and *Bombella*), we combined conditioned media assays with genomic and metabolomic analyses to investigate the basis of fungal growth modulation. We found that bacterial effects on fungi are mediated by secreted products, including medium acidification, with additional acidity-independent factors, resulting in selective inhibition of transient and potentially pathogenic fungi. Genomic and metabolomic analyses further identify candidate functions and compounds that may contribute to these effects.

## Results

### Lactobacillaceae and Acetobacteriaceae represent abundant and rare taxa sharing the larval diet with fungi

Three bacterial strains previously isolated from the *Scaptotrigona depilis* larval diet were used in this study (16). To provide eco-evolutionary context for these strains—phylogenetic placement, prevalence, and distribution within the bee colony—we sequenced their genomes using long- and short-read technologies to generate hybrid assemblies. We recovered high-quality genomes for all three strains (>98% completeness and <1% contamination; **Table S1**). Core genome phylogenetic analyses confirmed their placement, previously inferred from 16S rRNA amplicon data. The *Weissella* and *Bombella* isolates represent novel species (**Fig. 1**). *Weissella* sp. clusters with *W. sagaensis* and *W. hellenica* (88% ANI), taxa typically associated with fermented foods and bees (18). *Bombella* sp. is most closely related to *B. pluederhausenensis* and *B. dulcis* (80% ANI), and other bee-associated species (19, 20). In contrast, the *Apilactobacillus* isolate corresponds to the stingless bee-associated species *A. kunkeei*, commonly present during bee developmental stages (21). Consistent with this, its sister clade comprises bacteria from the genera *Acetilactobacillus* and *Nicoliella*, including strains recovered as metagenome-assembled genomes (MAGs) from the same larval diet samples (16).

**Fig. 1.**
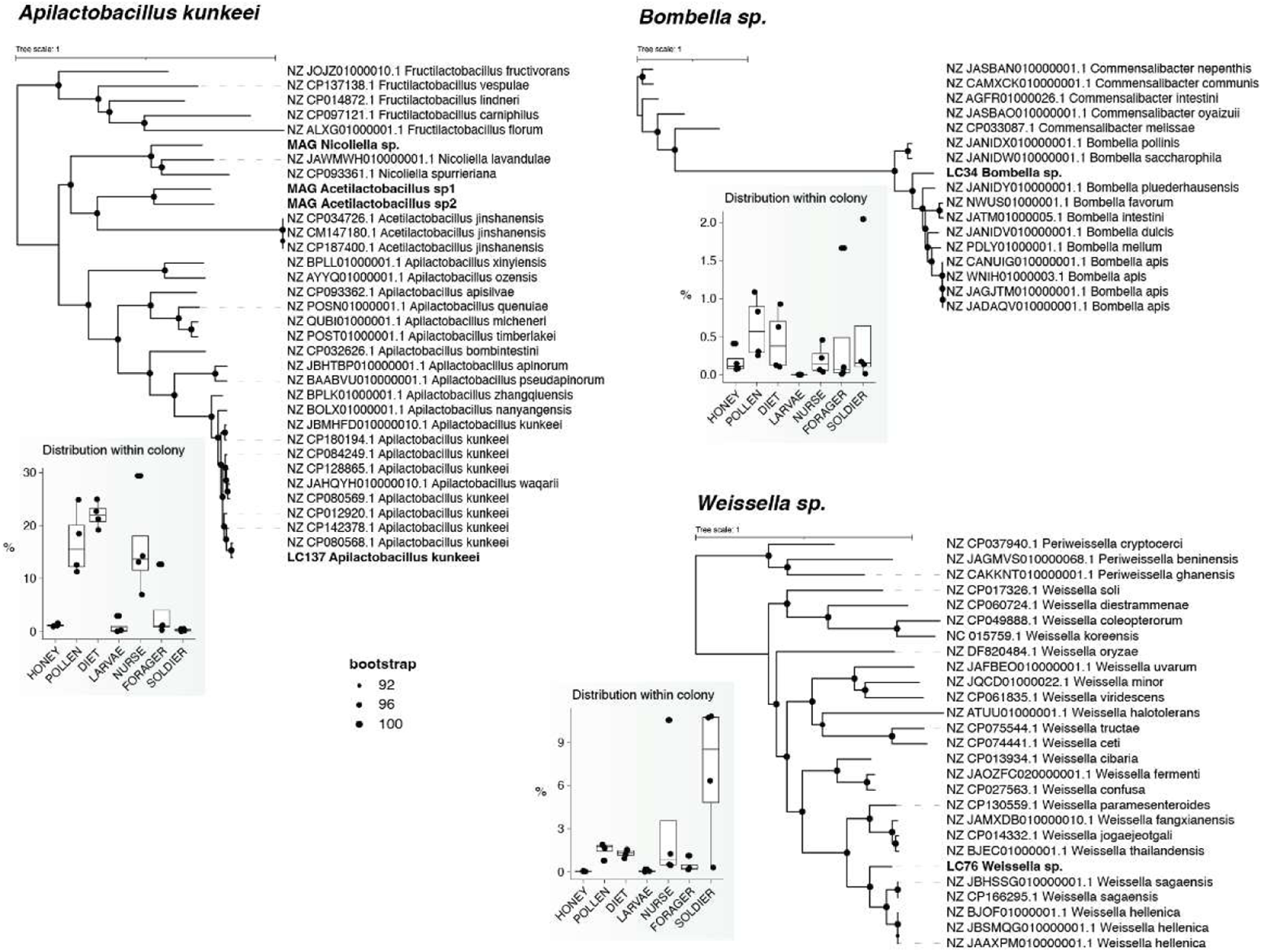
Taxonomy and distribution of three bacterial strains within stingless bee colonies. Maximum-likelihood phylogenies inferred from concatenated alignments of *Apilactobacillus* (377 single-copy orthologous genes; GTR+F+I+R6 substitution model; rooted using *Fructilactobacillus*), *Bombella* (965 single-copy orthologous genes; SYM+I+R5 substitution model; rooted using *Commensalibacter*), and *Weissella* (510 single-copy orthologous genes; GTR+F+I+R7 substitution model; rooted using *Periweissella*). Strains from this study are highlighted in bold, including MAGs from Caesar et al. 2025. Node support is indicated by filled circles (all bootstrap values >92%), and scale bars indicate substitutions per site. Box plots below each phylogeny show the relative abundance of each taxon across stingless bee colony components, including food stores (honey, pollen), brood (larval diet, larvae), and adult bees (nurse, forager, soldier).

To assess the distribution of the three bacterial species across hive components, we mapped metagenomic short reads from Caesar et al. (16) against a custom database comprising MAGs from that study and the genomes annotated here (see Materials and Methods). Although all three strains were isolated from the larval diet, read recruitment revealed that *Apilactobacillus* was the most abundant, particularly in the larval diet (∼23% of the bacterial community; **Fig. 1**). It was also prevalent in pollen and nurse bees, which provision the larval diet. *Weissella* showed lower overall abundance but was consistently detected across hive compartments, with higher proportions in soldier bees (up to 10%). *Bombella* was the least abundant, comprising <1% of the bacterial community in most samples, although slightly more prevalent in the larval diet and pollen.

### Bacteria-conditioned media inhibits pathogenic filamentous fungi

Competition assays with the three bacterial strains previously showed inhibition of filamentous fungi, including known bee pathogens (16). To test whether constitutively secreted products contribute to this effect, we quantified fungal growth (OD600) in the presence of bacteria-conditioned media (*i.e*., cell-free supernatants containing metabolites, signaling molecules, proteins, and other secreted factors). The fungi tested included one known bee pathogen (*Aspergillus* sp.), two species isolated from stingless bee brood (SB; *Talaromyces* sp. and *Penicillium* sp.), and two from honey bee brood (HB; *Mucor* sp. and *Penicillium* sp.).

Growth of *Aspergillus* was inhibited by conditioned media from all three bacterial species (one-way ANOVA, F₇,₁₆ = 45.12, p = 2.32 × 10⁻⁹; **Fig. 2, Table S2**). *Apilactobacillus* and *Weissella* showed the strongest effects, reducing growth by 74% ± 2% and 88% ± 0.7% (AUC, area under the curve). Inhibition was dose-dependent, with lower concentrations reducing growth by 27% ± 5% and 34% ± 4%. In addition to overall growth, spore production was quantified as a proxy for dispersal, although it did not consistently correlate with growth inhibition. For example, even low concentrations of conditioned media from both Lactobacillaceae members strongly suppressed sporulation, with effects comparable to high concentrations as for *Weissella* or exceeding as for *Apilactobacillus* (Welch’s ANOVA, F₇,₁₆ = 172.4, p = 7.42 × 10⁻¹⁴; **Table S2**).

**Fig. 2.**
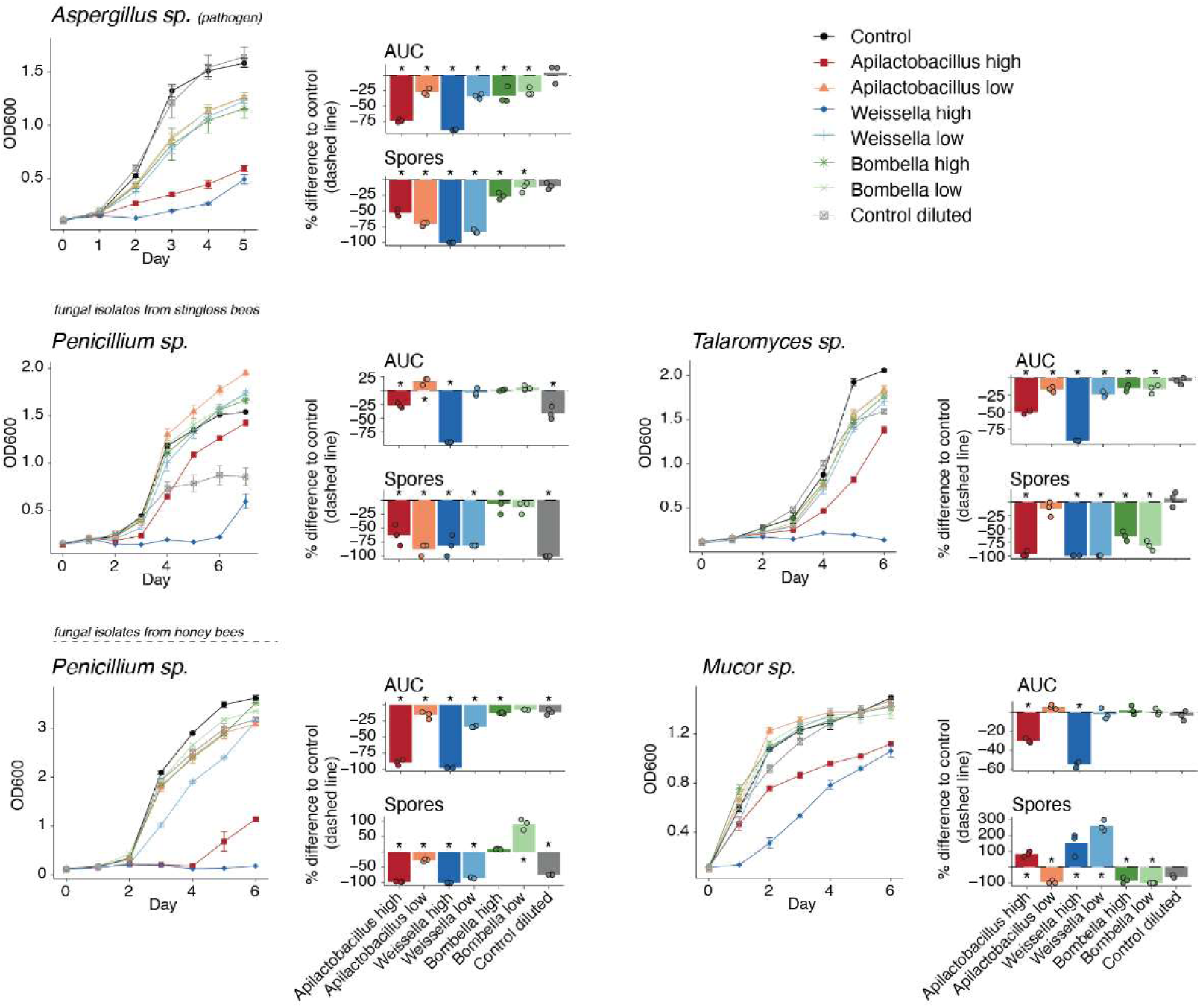
Effects of bacteria-conditioned media on filamentous fungal growth and sporulation. Each fungal species was grown in the presence of high (dark colors) and low (light colors) concentrations of bacteria-conditioned media (i.e., cell-free supernatants containing secreted metabolites and other factors). Controls included growth in the absence of conditioned media (black) and in medium diluted with sterile water to the same proportion as the high-dose treatment (gray). Left panels show growth curves over time, monitored until control reached a plateau or full sporulation. Right panels show the percentage change in area under the curve (AUC; total biomass) and spore production relative to the control (dashed line). Growth of all fungi was significantly affected by different treatments based on one-way ANOVA (or Welch’s ANOVA when variances were unequal). Asterisks (*) indicate significant differences from the control based on Dunnett’s multiple comparisons test performed on absolute values (adjusted p < 0.05). Exact p-values are provided in Table S2.

Across the other filamentous fungi, independent of bee species isolation source, *Apilactobacillus* and *Weissella* consistently showed stronger inhibition than *Bombella*, with *Weissella* generally having the greatest effect (**Fig. 2**; **Table S2**). At high concentrations, *Apilactobacillus* reduced growth by 27% ± 4% (stingless bee *Penicillium*) to 89% ± 4% (honey bee *Penicillium*), whereas *Weissella* caused 54% ± 3% (*Mucor*) to 97% ± 0.1% (honey bee *Penicillium*) inhibition. Even in cases of limited growth inhibition (e.g., stingless bee *Penicillium* with *Apilactobacillus*), conditioned media still strongly reduced sporulation (Welch’s ANOVA, F₇,₁₆ = 28.56, p = 6.81 × 10⁻⁸). Again, growth inhibition and sporulation were not always correlated. Although *Weissella* inhibited *Mucor* growth, the remaining mycelium showed increased sporulation relative to the control, while *Bombella* weakly inhibited *Talaromyces* growth but reduced spore production (**Fig. 2**). A dilution control confirmed that these effects were not explained by nutrient depletion: only *Penicillium* strains showed minor sensitivity to media dilution, with effects on growth and sporulation weaker than those of conditioned media (**Fig. 2**).

### Conditioned media inhibits non-symbiotic yeasts while promoting native *Zygosaccharomyces*

Given that bacteria-conditioned media inhibited filamentous fungi, we next asked whether it also affects yeast growth, which represents a core component of the stingless bee microbiome (16, 22). We included in our assays core mycobiome members isolated from stingless bee brood (SB; *Zygosaccharomyces* sp. and *Starmerella* sp.), as well as corresponding species isolated from honey bee brood (HB).

Conditioned media affected yeast growth, although effects were weaker and more variable than for filamentous fungi, with no near-complete inhibition (**Fig. 3**). *Apilactobacillus* and *Bombella* conditioned media consistently reduced yeast growth, with up to 29% ± 5% reduction in AUC for honey bee *Zygosaccharomyces* (Welch’s ANOVA, F₇,₁₆ = 4.38, p = 0.00691) *Starmerella* isolates were more broadly sensitive, showing dose-dependent inhibition by conditioned media from all strains (one-way ANOVA, HB, F₇,₁₆ = 66.41, p = 1.24 × 10⁻¹⁰; Welch’s ANOVA, SB, F₇,₁₆ = 7.70, p = 3.79 × 10⁻⁴). In contrast, stingless bee *Zygosaccharomyces* was not inhibited; *Apilactobacillus* and *Weissella* increased its growth up to nearly two-fold at high concentrations (one-way ANOVA, F₇,₁₆ = 5.95, p = 0.00155). For the stingless bee *Starmerella* and honey bee *Zygosaccharomyces*, however, the dilution control indicated that inhibition may partly reflect reduced nutrient availability (**Fig. 3**). Indeed, the strongest inhibition of honey bee *Zygosaccharomyces*—by *Bombella*—fell within the dilution-control range; however, high concentrations of *Weissella* conditioned media, which similarly dilute the medium, did not reduce growth, suggesting that nutrient depletion alone does not explain the inhibitory effects.

**Fig. 3.**
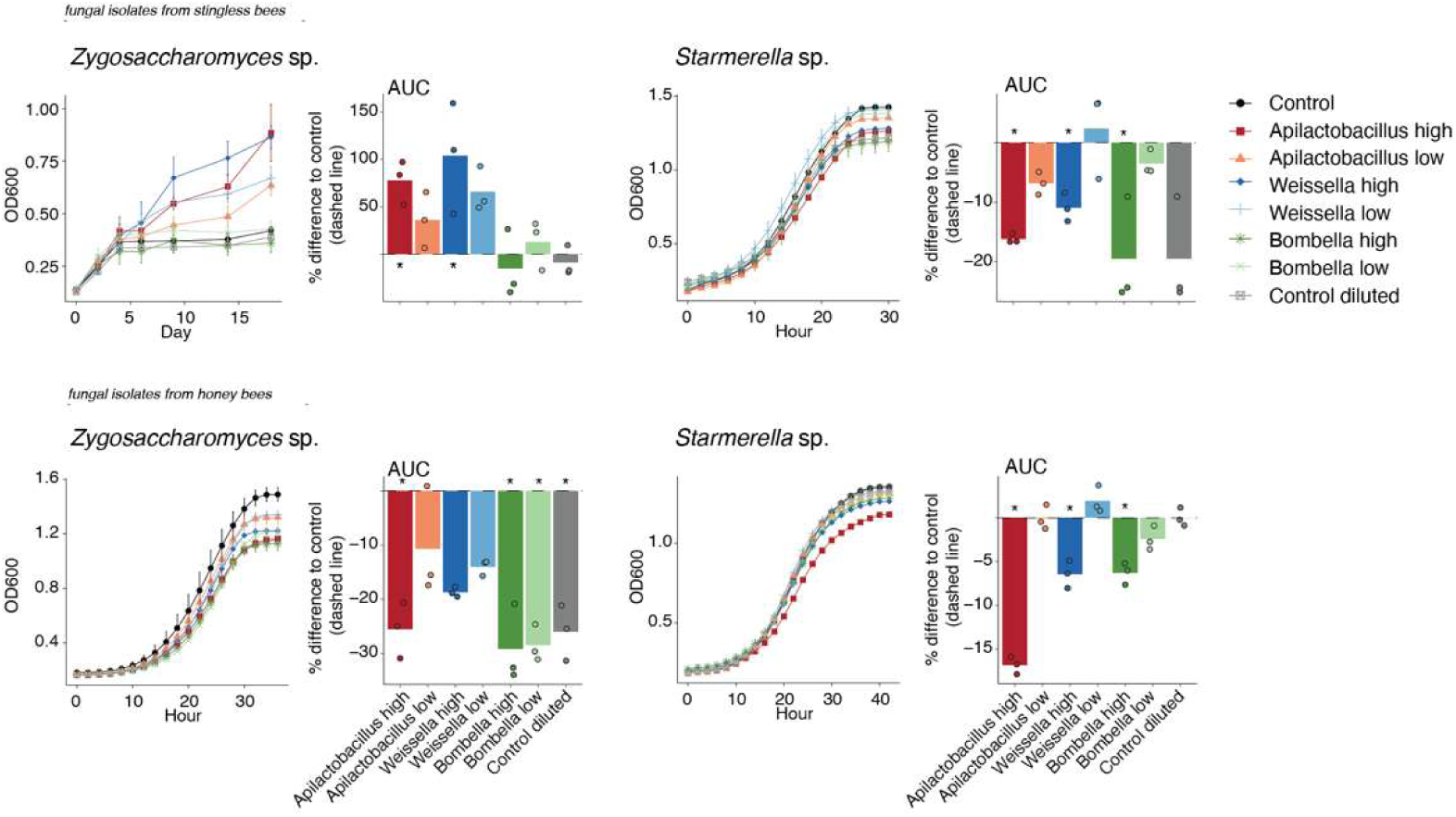
Effects of bacteria-conditioned media on yeast growth. Each yeast species was grown in the presence of high (dark colors) and low (light colors) concentrations of bacteria-conditioned media. Controls included growth in the absence of conditioned media (black) and in medium diluted with sterile water to the same proportion as the high-dose treatment (gray). Left panels show growth curves over time, monitored until control reached a plateau or full sporulation. Right panels show the percentage change in area under the curve (AUC; total biomass) relative to the control (dashed line). Growth of all fungi was significantly affected by different treatments based on one-way ANOVA (or Welch’s ANOVA when variances were unequal). Asterisks (*) indicate significant differences from the control based on Dunnett’s multiple comparisons test performed on absolute values (adjusted p < 0.05). Exact p-values are provided in Table S2.

### Effects of conditioned media on fungal growth go beyond media acidification

The lactic acid bacteria *Apilactobacillus* and *Weissella* showed the strongest inhibitory effects overall (**Fig. 2**, **Fig. 3**). Because these bacteria can acidify the medium to pH 4.2, and low pH was shown to inhibit the filamentous fungi in agar (16), we tested whether acidification alone could explain the effects of conditioned media. Conditioned media were adjusted to pH 6 prior to fungal assays to test whether inhibition persisted. In parallel, low-pH controls, including lactic acid, tested whether low pH alone recapitulated the inhibition. Effects retained after conditioned media pH adjustment indicate a role for bacterial secreted products; effects observed only at low pH controls suggest a pH-driven mechanism; and those observed under both conditions suggest combined effects of acidity and other bacterial products.

Both *Apilactobacillus* and *Weissella* conditioned media adjusted to near-neutral pH still inhibited the pathogen *Aspergillus* (one-way ANOVA, F₄,₁₀ = 44.87, p = 2.35 × 10⁻⁶), reducing growth by 56% ± 3% and 58% ± 2% (**Fig. 4, Table S2**). However, both low-pH controls also reduced *Aspergillus* growth, indicating that acidification contributes to these effects (16). Consistently, non-adjusted conditioned media (low pH; **Fig. 2**) showed stronger inhibition than pH-adjusted treatments (**Fig. 4**). Across other filamentous fungi, low-pH controls also reduced growth (**Fig. S1, Table S2**), while near-neutral conditioned media retained inhibitory activity in most cases. Yeasts showed more variable responses. Neither pH-adjusted conditioned media nor low pH consistently promoted the stingless bee-associated *Zygosaccharomyces* (Welch’s ANOVA, F₄,₁₀ = 1.79, p = 0.208; **Fig. 4)**, suggesting that the previously observed growth promotion depends on combined effects of bacterial metabolites and acidification, as for most filamentous fungi. In contrast, the non-symbiotic honey bee *Zygosaccharomyces* remained inhibited by pH-adjusted *Apilactobacillus* conditioned media (Welch’s ANOVA, F₄,₄.₇₄ = 73.44, p = 0.0001; 53% ± 4%), with no inhibition in the pH controls. *Apilactobacillus* also inhibited *Starmerella* sp. (**Fig. S1, Table S2**), although less than the non-symbiotic *Zygosaccharomyces*. *Weissella* similarly inhibited *Starmerella*, whereas only the stingless bee-associated *Starmerella* was sensitive to the low-pH (HCl) control.

**Fig. 4.**
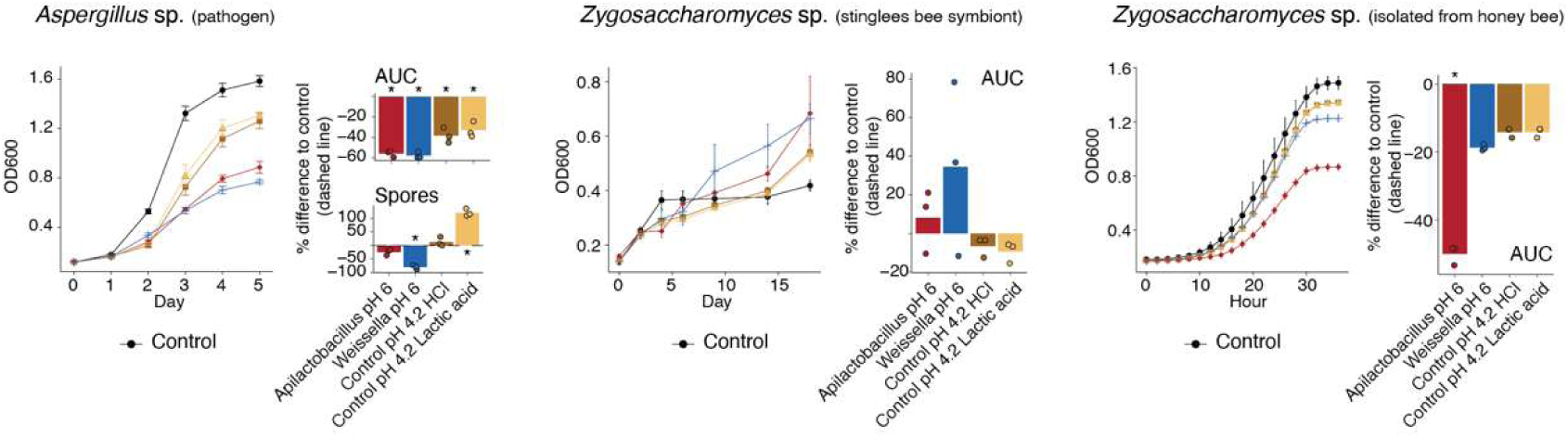
Effects of pH on fungal growth. Each fungal species was grown in the presence of high concentrations of bacteria-conditioned media adjusted to pH 6 (red and blue). Controls included fresh medium without conditioned media (black, dashed line) and fresh medium adjusted to pH 4.2 with HCl or lactic acid (brown and yellow). Left panels show growth curves over time, monitored until control cultures reached a growth plateau or full sporulation. Right panels show the percentage change in area under the curve (AUC; total biomass) relative to the control (dashed line) and, for *Aspergillus*, spore production. Growth of *Aspergillus* and honey bee *Zygosaccharomyces* was significantly affected by different treatments based on one-way ANOVA (or Welch’s ANOVA when variances were unequal). Asterisks (*) indicate significant differences from the control based on Dunnett’s multiple comparisons test performed on absolute values (adjusted p < 0.05). Exact p-values are provided in Table S2.

### Stingless bee-associated bacteria encode and produce candidate molecules involved in microbial interactions

To identify bacterial factors potentially underlying the observed effects on fungal growth, we combined genome-resolved annotation with untargeted metabolomics. Genomes of the *A. kunkeei*, *Weissella* sp., and *Bombella* sp. were screened for genes associated with the production and secretion of bioactive compounds, including secondary metabolites, antimicrobial peptides, and carbohydrate-active enzymes. In parallel, we performed untargeted LC-MS/MS profiling of bacterial cultures to characterize the secreted metabolome and identify candidate small molecules.

Genome annotation of *Apilactobacillus* and *Weissella* identified features consistent with indirect fungal interactions (**Fig. 5A**), including L- and D-lactate dehydrogenases (*ldh*, *ldhD*) supporting acidification, and stress- and pH-homeostasis functions such as HSP20-family proteins and Na+/H+ antiporters (*napA*). Predicted secreted proteins included OppA and the serine protease HtrA, while only *Weissella* encoded a secreted fungal cell wall-targeting GH18 chitinase. Secondary metabolism analysis identified a shared terpene precursor pathway and T3PKS locus, while *Weissella* additionally encoded a predicted RiPP cluster. The T3PKS locus is conserved across the *Apilactobacillus-Acetilactobacillus-Nicoliella* clade but lacks a canonical type III polyketide synthase and associated tailoring genes, instead containing primary metabolic genes (e.g., HMG-CoA synthase), suggesting a conserved metabolic rather than secondary metabolite locus (23). In contrast, *Bombella*—which has a lower prevalence and weaker fungal inhibition—encodes a broader repertoire of biosynthetic gene clusters, similar to *B. apis* from honey bees (2), including deazapurine, terpene, T1PKS, redox-cofactor, and RiPP (linocin) clusters (**Fig. 5A, Table S3**).

**Fig. 5.**
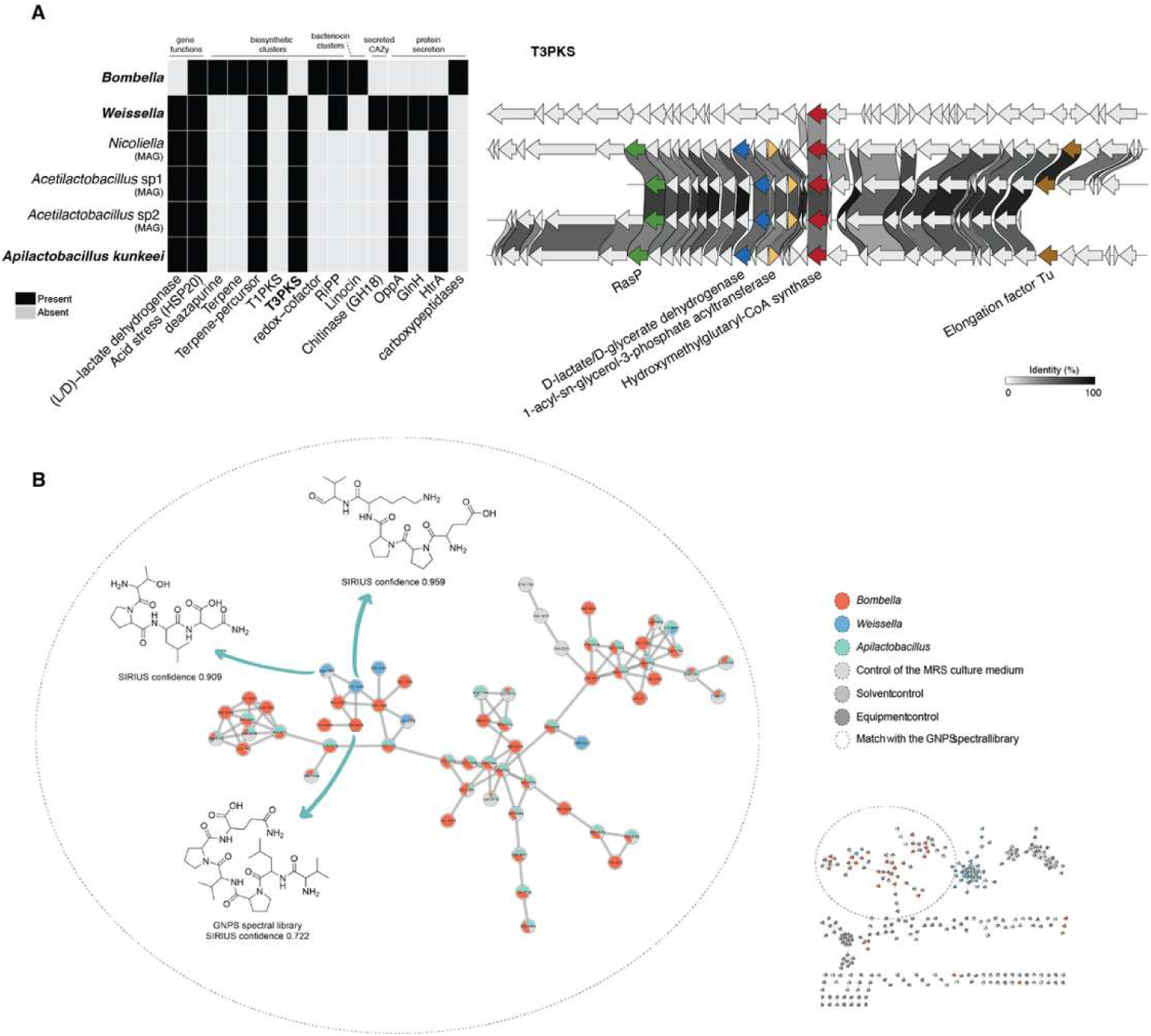
Genomic and metabolomic characterization of stingless bee-associated bacteria. (**A**) Left, presence (black) and absence (light gray) of selected genomic features potentially associated with fungal interactions across bacterial strains (see Table S3 for full annotation). Results are shown for the three bacterial isolates, as well as the three metagenome-assembled genomes (MAGs) representing the most abundant taxa in the larval diet and closely related to *Apilactobacillus*. Features include general gene functions (Prokka, eggNOG), biosynthetic gene clusters (antiSMASH), bacteriocin clusters (BAGEL4), secreted CAZymes (dbCAN + SignalP), and other secreted proteins (SignalP). Right, zoom-in of T3PKS-associated genomic regions across representative Lactobacillaceae isolates, including MAGs of abundant strains in stingless bee larval diet. Arrows indicate predicted coding sequences; highlighted genes correspond to core and additional biosynthetic genes, and shading between CDS indicates protein identity. (**B**) Molecular network generated from untargeted LC-MS/MS data using feature-based molecular networking. Nodes represent molecular features, and edges indicate spectral similarity. Nodes are colored by bacterial source or controls. A highlighted cluster (dashed outline) contains peptide-like molecules annotated based on GNPS spectral library matches and SIRIUS predictions (confidence values indicated). Insets show the full network and the position of the highlighted cluster.

Untargeted LC–MS/MS and feature-based molecular network analyses revealed a peptide-enriched molecular family predominantly composed of metabolites shared among *Bombella*, *Weissella*, and *Apilactobacillus* (**Fig. 5B**). This molecular family was mainly formed by nodes derived from bacterial cultures, although a subset of signals was also detected in the MRS medium control, indicating a possible contribution from peptide components naturally present in the culture medium. Nevertheless, several nodes were exclusive to or more abundant in bacterial cultures, suggesting bacterial transformation and/or production of related peptide derivatives. *In silico* annotation using GNPS and SIRIUS indicated that compounds within this molecular family correspond mainly to small oligopeptides, including pentapeptides, with some predicted structures showing high confidence scores (SIRIUS confidence up to 0.959). The observed profiles suggest that these bacteria share metabolic capabilities related to the processing, modification, or secretion of small peptides, consistent with the metabolic behavior typically associated with fermentative bacteria inhabiting nutrient-rich environments (24, 25)

## Discussion

Microbial interactions are a key driver of microbiome assembly and can act through both direct mechanisms, such as the production of antimicrobial compounds, and indirect mechanisms, such as environmental modification (e.g., acidification) (14, 26, 27). In stingless bees, we previously showed (16) that the larval diet hosts a stable multi-kingdom community in which bacteria inhibit potentially pathogenic filamentous fungi while allowing the persistence of beneficial yeast symbiont. Here, we show that these effects on fungal growth are mediated by bacterial secreted products, including organic acids that lower substrate pH, as well as additional acidity-independent compounds. In some interactions, either mechanism alone is sufficient to inhibit fungal proliferation, whereas in others their combined effects result in stronger inhibition or, in the case of the symbiotic yeast, promotion.

The stingless bee larval diet consists of a mass-provisioned mixture of pollen, nectar, and bee secretions, forming a sugar-rich and acidic substrate (28). Host-derived compounds and enzyme-driven processes can contribute to this acidity, as observed in the nurse-produced larval and queen diet of honey bees (royal jelly) (29). However, because microbes are present in the larval diet from the time it is deposited in brood cell (16), the relative contributions of host secretions and microbial metabolism (bacterial and yeast-derived) to environmental acidification cannot be readily disentangled. Metagenomic analyses and the distribution patterns of the isolates used in this study confirm that Lactobacillaceae dominate the larval diet microbiome, particularly members of the *Apilactobacillus-Acetilactobacillus-Nicoliella* clade. As expected for lactic acid bacteria, and supported by both *in vitro* phenotyping (16) and genomic annotation, these taxa actively acidify the larval diet.

The high-dose conditioned media from *Apilactobacillus* and *Weissella*, with pH ∼4.2, showed the strongest and broadest inhibitory effects on fungal isolates, particularly filamentous fungi, including the bee pathogen *Aspergillus*—consistent with the competition assays from our previous study (16). This inhibitory effect was partially recovered by pH controls, as media acidified with either HCl or lactic acid also inhibited filamentous fungi across species. Together, these observations indicate that bacteria-driven substrate acidification contributes to fungal inhibition. This finding is consistent with the well-established role of organic acid-producing bacteria in restricting fungal growth across diverse systems, including human microbiomes, insect models, and food fermentation (30–36). However, acidification alone does not fully explain the observed patterns. In several cases, inhibition of potentially pathogenic fungi was stronger in conditioned media than in acidified controls and persisted after increase in pH, indicating that additional acidity-independent mechanisms contribute to fungal inhibition.

The persistence of inhibition after increase in pH suggests that bacterial secondary metabolites, *i.e.,* secreted products other than organic acids, contribute to fungal inhibition. Lactobacillaceae are known to produce secondary metabolites with antifungal activity. For example, *Lactobacillus plantarum* secretes metabolites such as 3-phenyllactic acid and benzeneacetic acid, 2-propenyl ester, which have been suggested to play a role in inhibiting plant pathogenic fungi, including *Penicillium* and *Fusarium* (37). Similarly, another *Lactobacillus* species produces the metabolite 1-acetyl-β-carboline, which inhibits *Candida albicans* hyphal morphogenesis and biofilm, processes required for its virulence (38). In the stingless bee-associated strains, however, canonical biosynthetic pathways typically associated with antifungal activity were not detected. Even antibacterial features previously reported in an *Apilactobacillus kunkeei* isolate from honey bees—the plasmid-encoded T1PKS and the lanthipeptide kunkecin A (39, 40)—were absent in the strain studied here, which do not carry a plasmid. In the stingless bee-associated *A. kunkeei* and related strains, predicted T3PKS and terpene-associated loci lacked key biosynthetic features (41–44) and showed no corresponding metabolite signals. In contrast, the most promising BGC identified was a T1PKS cluster encoded by *Bombella*, which, although not highly prevalent across stingless bee hive microbiome, is consistent with previous findings linking *Bombella apis* to antifungal activity in honey bee colonies (2). The cluster encodes characteristic PKS core and reductive tailoring domains, but shows low similarity to characterized BGCs (MIBiG similarity score of 0.66 to the 6-methylsalicylic acid cluster from *Aspergillus terreus*), suggesting the potential production of a chemically novel bioactive metabolite. Complementary metabolomic analyses revealed extracellular peptide-like compounds across all strains, including *A. kunkeei*, while genomic analyses identified numerous SignalP-positive hypothetical proteins. Although these predictions likely include a mixture of ordinary secreted, membrane-associated, and cell-wall-associated proteins, they nonetheless point to a substantial set of exported proteins with currently unknown functions. Together, these results suggest that fungal inhibition by stingless bee-associated lactic acid bacteria may involve both canonical secondary metabolite pathways and non-canonical secreted compounds that remain undetected by current genome-mining approaches. Consistent with this, variation in fungal responses—hyphal inhibition in some cases and reduced sporulation in others—indicates multiple mechanisms targeting different stages (26, 45, 46).

In contrast to filamentous fungi, yeast responses to bacterial secreted products were more variable and, in some cases, contrasting. Acid-tolerant yeasts, such as those from the *Zygosaccharomyces* genus, are well adapted to maintain intracellular pH homeostasis and resist weak acid stress through efficient proton extrusion and organic acid detoxification systems (47–49). Consistent with this, the non-symbiotic *Zygosaccharomyces* isolate was not strongly inhibited by substrate acidification alone, but was inhibited by conditioned media after increase in pH. In contrast, the symbiotic *Zygosaccharomyces* from stingless bees was promoted in conditioned media at acidic pH, likely reflecting species-specific differences in tolerance to bacterial products and environmental conditions, and potentially consistent with its co-diversification with the bee host (22). Beyond resistance to bacterial products, additional interactions may contribute to the observed promotion of the symbiotic yeast, potentially including metabolic interactions such as cross-feeding, as described in other multi-kingdom microbiomes (50–52).

By integrating functional assays with genomic and metabolomic analyses, we identified candidate mechanisms underlying bacterial-fungal interactions in bees. Although no specific compound could be linked to the observed fungal growth responses, the combined data point to fermentation, extracellular peptide processing, and, in *Weissella*, chitin degradation as potential mechanisms. The concentration-dependent effects of conditioned media further suggest that these interactions depend on the accumulation of bacterial metabolites or bacterial-induced changes to the extracellular chemical environment (53). Together, these findings provide potential mechanisms through which changes in microbiome composition or activity could influence fungal growth and bee health.

## Material and Methods

### Microbial strains and cultivation for conditioned media assay

Bacterial and fungal strains isolated from stingless bee brood (SB; *Scaptotrigona depilis,* SISGEN permission no. AE1AFA6) and honey bee brood (HB; *Apis mellifera*) were used for conditioned media assays. Bacterial strains were grown in 5 mL MRS broth for 24 h at 250 rpm and 30°C (stingless bee isolates) or 34°C (honey bee isolates) (16, 54, 55). Cultures were centrifuged at 5,000 rpm for 8 min, and supernatants containing bacterial byproducts were filtered through sterile 0.22-μm Steriflip tubes to obtain conditioned media. Filamentous fungi were grown on MRS agar plates at their respective temperatures. After sporulation, spores were harvested by adding 2 mL PBS 1X to the plates and scraping the surface. Spore concentrations were adjusted to 50 spores/μL, and 2 μL was added to each well (100 spores/well). Yeasts were grown on MRS agar plates at their respective temperatures for 48 h, harvested with a sterile stick, and diluted in PBS 1X to an OD600 of 1.0. Bacterial and fungal cultures were initiated in advance to allow collection of conditioned media, yeast cells, and filamentous fungal spores on the day the experiment began.

### Conditioned-media assay

The effects of conditioned media (CM) from three bacterial strains on fungal growth were tested in 96-well plates with three replicates per condition. Each well contained 200 µL of medium plus 2 µL fungal inoculum (prepared as described above). Conditions were: 200 µL MRS (Control); 100 µL MRS + 100 µL CM (High); 175 µL MRS + 25 µL CM (Low); and 100 µL MRS + 100 µL sterile water to control for nutrient dilution (Control diluted). To distinguish pH effects from those of bacterial metabolites, CM from Lactobacillaceae strains, which reduce medium pH to ∼4.2 (16), was adjusted to pH 6 with 10 µL of 10 mM Tris-HCl and mixed with 100 µL MRS (pH 6). Additional pH controls without bacterial metabolites consisted of MRS adjusted to pH 4.2 with either 1.2 µL of 10 mM HCl (Control pH 4.2 HCl) or 2 µL DL-lactic acid (∼14 g L⁻¹; Control pH 4.2 Lactic acid). Plates were incubated at the respective fungal growth temperatures, corresponding to the hive temperatures of their bee hosts. Fungal growth was monitored by optical density at 600 nm (OD600). For filamentous fungi, OD600 was measured daily until sporulation occurred in the positive control, after which each well was diluted with 200 µL PBS and spores were counted using a C-chip hemocytometer. For yeasts, OD600 was measured at 2-h intervals until the control reached a growth plateau (∼35 h), except for the more slowly growing *Zygosaccharomyces* symbiont of stingless bees, which was measured at longer intervals. All plates included negative controls containing 100 µL fresh medium and 100 µL sterile medium without fungal inoculum, for which no growth was expected.

### Growth curve analyses

Growth curves were visualized in R using ggplot2 (56) and fitted for each replicate using GrowthCurver (57), which models microbial growth with a logistic equation. The area under the fitted logistic curve (auc_l = AUC) was extracted for each replicate as a summary metric integrating growth rate and final biomass. Replicate AUC values were analyzed in R (**Table S2**) using one-way ANOVA or Welch’s ANOVA when homogeneity of variances was not met, as determined by Bartlett’s test, with condition as the explanatory factor. Treatments were compared with the control using Dunnett’s test with adjustment for multiple comparisons (α = 0.05). For visualization, the percentage change in AUC relative to the control was calculated for each replicate and plotted using ggplot2 (56). The same statistical approach was applied to filamentous fungal spore counts, which were visualized as percentage differences relative to the control.

### DNA extractions and genome sequencing

DNA was extracted using the ZymoBIOMICS™ DNA Miniprep Kit following the manufacturer’s protocol and quantified using a Qubit fluorometer. Short-read libraries were prepared with the Illumina DNA Prep kit and IDT 10 bp UDI indices and sequenced on an Illumina NextSeq 2000 (2 × 151 bp). Demultiplexing, quality control, and adapter trimming were performed with bcl-convert (v3.9.3). For long-read sequencing, libraries were prepared using the Oxford Nanopore Technologies Ligation Sequencing Kit (SQK-NBD114.24) with the NEBNext Companion Module (E7180L) without additional fragmentation or size selection. Sequencing was performed on MinION Mk1B or GridION devices using R10.4.1 flow cells (400 bp/s mode, minimum read length 200 bp). Basecalling, demultiplexing, and adapter trimming were performed with Guppy (v6.4.6, super-accurate model).

### Genome assembly and phylogeny

Illumina reads were quality-trimmed using Trimmomatic v0.36 (LEADING:28 TRAILING:28 MINLEN:75) (58), and Nanopore reads were filtered with Filtlong (--min_length 1000, --keep_percent 95, --target_bases 400000000) (59). Filtered short and long reads were hybrid assembled using Unicycler v0.5.0 with default parameters (60). Assembly quality metrics and genome statistics were assessed with QUAST v5.3.0 (61). Genome completeness and contamination were estimated using CheckM v1.2.2 (62). To estimate genome coverage, Illumina reads were mapped to the assemblies using Bowtie2 v2.5.1 (63), and coverage statistics were calculated with SAMtools v1.17 (64). Gene prediction and initial annotation were performed using Prokka v1.14.6 with default settings (65). Predicted proteins were queried against the NCBI nr database using DIAMOND blastp (--max-target-seqs 1 --outfmt 6) to guide genome selection for phylogenetic analysis (66). Single-copy orthologs were identified using OrthoFinder v3.0.1b1 with default parameters (67). Orthologous genes were aligned with MAFFT (--globalpair --maxiterate 1000) and trimmed using trimAl v1.5.rev0 (-automated1) (68, 69). Trimmed alignments were concatenated and used for phylogenetic inference with IQ-TREE v2.3.6 (--seqtype DNA -m MFP -B 1000) (70). Trees were visualized using iTOL (71).

### Within-hive distribution of the three bacterial strains

Illumina shotgun metagenomic reads available from the NCBI Sequence Read Archive (SRA; BioProject PRJNA1216660) were used to estimate the relative abundance of each bacterial strain across colony components, including honey pots, pollen pots, larval diet, larvae, nurse bees, forager bees, and soldier bees. The genomes of the three isolates were incorporated into a reference database together with metagenome-assembled genomes (MAGs) recovered from the same study. Inclusion of MAGs enabled competitive mapping and reduced bias associated with mapping reads to isolate genomes alone. Illumina reads were mapped to the reference assemblies using Bowtie2 v2.5.1 (--no-discordant --no-mixed) (63), and mapping statistics were obtained using SAMtools v1.17 idxstats (64). Relative abundance was estimated as the proportion of reads mapping to each genome and visualized using ggplot2 (56).

### Prediction of biosynthetic gene clusters and secreted enzymes

Genome annotations generated with Prokka v1.14.6 were used for downstream analyses (65). Functional annotation was complemented using eggNOG-mapper v2.1.12 (-m diamond) (72). Biosynthetic gene clusters were predicted using antiSMASH v8.0.4 with comprehensive detection modules enabled (73). Bacteriocin and antimicrobial peptide clusters were identified using BAGEL4 via the web server (74). Carbohydrate-active enzymes (CAZymes) were predicted using dbCAN v5.2.8 (75), and secretion signals were identified using SignalP 6 (--mode fast, --organism other) to predict potentially secreted proteins (76). These analyses were also performed on three metagenome-assembled genomes (MAGs) representing the most abundant bacterial taxa in the larval diet for comparison (16).

### Metabolite Extraction and UHPLC–MS/MS-Based Metabolomics

Bacterial cultures were reactivated from MRS glycerol stocks by inoculating 100 μL into 5 mL MRS broth for 24 h at 30 °C and 150 rpm. Pre-cultures (1 mL) were then transferred to 20 mL MRS broth and incubated under the same conditions for 24 h. For large-scale cultivation, 5 mL of culture were inoculated into 250 mL Erlenmeyer flasks containing MRS broth (in triplicate) and incubated at 30 °C and 150 rpm. After 24 h, Diaion HP-20® resin (70 g/L) was added and recovered three days later for extraction with 100% acetone. Extracts were fractionated using C18 solid-phase extraction (SPE) cartridges (Discovery DSC-18®, Supelco) with 25%, 50%, and 100% MeOH/H2O. The 25% MeOH fractions were discarded, and the 50% and 100% fractions were combined, concentrated, resuspended, and analyzed at 100 ppm.

Metabolomic analyses were performed by UHPLC-ESI-MS/MS using an Agilent 6545 ESI-QTOF-MS coupled to an Agilent 1290 Infinity II UHPLC system (Agilent Technologies, Santa Clara, CA, USA) at the Central Analytical Facility, Department of Chemistry, Federal University of São Carlos (UFSCar). Data were acquired in positive ionization mode with MS and MS/MS scan ranges of *m/z* 150–1500 and 70–1500, respectively, at 3 spectra/s. Chromatographic separation used a Zorbax EC-C18 column (1.8 μm, 4.6 × 50 mm) with H2O + 0.1% formic acid (A) and ACN + 0.1% formic acid (B) as mobile phases. Collision-induced dissociation (CID) spectra were acquired using a collision energy ramp. Raw data were converted to .mzXML using MSConvert and processed in MZmine 4.7.8. Feature-based molecular networking (FBMN) was performed on GNPS2 (77, 78) using precursor and fragment ion tolerances of 0.08 and 0.1 Da, respectively. Networks were generated with cosine scores >0.65 and ≥4 matched fragment ions, retaining only molecular families with ≥2 nodes; GNPS2 spectral library matches were filtered using the same criteria. Networks were visualized in Cytoscape 3.10.2, and MS/MS data were additionally analyzed in SIRIUS for molecular formula prediction and *in silico* compound annotation (79, 80).

## Acknowledgments

We thank the Newton Lab members at Indiana University for their valuable feedback throughout this study and for helping set up strains for growth when needed to coordinate experiments. LC and IN acknowledge funding support from the National Science Foundation (NSF) IOS Award #2005306 and the GEMS Biology Integration Institute, funded by the NSF DBI Biology Integration Institutes Program, Award #2022049. GT and MTP acknowledge the financial support of São Paulo Research Foundation (FAPESP) grant #2025/10223-4 (MTP), and Conselho Nacional de Pesquisa e Desenvolvimento Tecnológico (CNPq) grants #142024/2020-1 (GTP) and #307893/2022-7 (MTP).

## Data availability

Genome data are available through the NCBI BioSample database under accession numbers SAMN63108970, SAMN63108971, and SAMN63108972.

